# Role of Membrane-tension Gated Ca Flux in Cell Mechanosensation

**DOI:** 10.1101/134395

**Authors:** Lijuan He, Jiaxiang Tao, Fangwei Si, Yi Wu, Tiffany Wu, Vishnu Prasath, Denis Wirtz, Sean X. Sun

## Abstract

Eukaryotic cells are sensitive to mechanical forces that they experience from the environment. The process of mechanosensation is complex, and involves elements such as the cytoskeleton and active contraction from myosin motors. Ultimately, mechanosensation is connected to changes in gene expression in the cell, or mechanotransduction. While the involvement of the cytoskeleton in mechanosensation is known, processes upstream to cytoskeletal changes is unclear. In this paper, using a microfluidic device that mechanically compresses live cells, we demonstrate that calcium currents and membrane tension-sensitive ion channels directly signals to the Rho GTPase and myosin contraction. In response to membrane tension changes, cell actively regulates cortical myosin contraction to balance external forces. The process is captured by a mechanochemical model where membrane tension, myosin contraction and the osmotic pressure difference between the cytoplasm and extracellular environment are connected by mechanical force-balance. Finally, to complete the picture of mechanotransduction, we find that the tension-sensitive transcription factor YAP translocates from the nucleus to the cytoplasm in response to mechanical compression.

## Introduction

Mechanotransduction, the conversion of physical force into biochemical information inside the cell, is a complex process that regulates a large variety of physiological processes such as embryogenesis and tissue growth [1, 2]. Dysregulation of mechanotransduction are implicated in the development of major human diseases, such as cancer and arteriosclerosis [3-7]. The most upstream process in mechanostranduction is mechanosensation: the earliest step in which the cell senses changes in the external mechanical environmental and/or forces. It is known that the actin cytoskeleton and myosin contractility are required for mechanosensation and mechanotransduction. Though there are many studies that examine how forces regulate cytoskeletal dynamics in vitro [8,9], precisely how the F-actin network and myosin are signaled in live cells by external forces during mechanosensation is unclear. In this paper, by mechanically compressing live cells, we identify transmembrane calcium currents and membrane tension-sensitive cation channels are responsible for activating RhoA GTPase, which regulates non-muscle myosin II assemblies in the cell cortex and cytoplasm. These experimental results, together with a mechanical model of the cell cortex, suggests that the cell maintains a homeostatic value of membrane tension, and activates myosin contraction in response to tension changes. This feedback loop leads to a dynamic adjustment of active stress generated by the cell, and ultimately can explain main features of mechanosensation.

Cortical tension and myosin contraction in tissue cells are biochemically controlled by the Rho family of small GTPases, especially RhoA [5]. RhoA switches between a GTP-bound, active state and a GDP-bound, inactive state, which signals to the Rho-associated kinase, ROCK. ROCK phosphorylates myosin light chain, which then controls mini-filament assembly and generation of active contractile stress. Externally applied mechanical forces can trigger the response of RhoA. For example, the active form of RhoA increases when cells are mechanically pulled by magnetic tweezers [10,11]. High shear stress (65 dyn/cm^2^) on bovine aortic endothelial cells leads to decrease in RhoA activity [12], whereas low shear stress induces an initial increase in RhoA activity, which is followed by returning to control levels after 10 minutes [13].

To investigate the mechanosensation in live cells in real time, we established a microfluidic-based mechanical compression system, in which the live cells can dynamically switch between confined and un-confined status. A FRET based sensor is used to monitor the real-time response of RhoA activity in cells when they are subjected to different environments. We find that the mechanical compression leads to an immediate drop in RhoA activity as indicated by the RhoA FRET sensor. The decreased RhoA activity is maintained while the cell is compressed. Upon decompression, RhoA activity resumes to the original level. Either depriving cells of calcium or blocking transient receptor potential cation channel subfamily V member 4 (TRPV4) significantly decreases the change in RhoA activity in response to the mechanical shock. Moreover, inhibiting myosin activity by blebbistatin does not affect RhoA activity change during compression. These results can be recapitulated in a computational mechanical model of cell mechanosensation where membrane and cortical tensions are explicitly connected to an externally applied force. Conceptually, the results and the model suggest that mechanosensation partly arises from a negative feedback control system that maintains a homeostatic membrane tension. To connect mechanosensation with downstream mechanotransduction, we further reveal that the Yes-associated protein (YAP) transcription factor largely left the nucleus and distributed more in the cytoplasm upon compression. This suggests that there is a direct link between physical forces, cell cortical tension and YAP transcriptional activity, as revealed by parallel studies in other settings [14,15].

Our results are relevant for understanding how cells respond to external mechanical forces, and interact with physically confined environments. When metastatic cancer cells leave their primary tumor sites and migrate form a metastasis tumor, they move within and between three-dimensional tissues, capillaries and lymph nodes, the properties of which cannot be fully recapitulated by 2D petri dishes [16-19]. Similarly, cells of the immune system, such as dendritic cells, also migrate within tissues to sample different environments [20]. Cells also experience mechanical forces from the surrounding matrix [21] as well as from other cells in different environments [22-25]. Our work identifies the most upstream signals that allow cells to respond to mechanical forces and physical changes. Identification of calcium as the primary signal is also consistent with observations in cells under mechanical stretch [26], and opens doors to manipulate mechanosensation properties of live cells.

## Materials and Methods

### Cell culture

Human fibrosarcoma HT1080 (ATCC) cells were cultured in Dulbecco’s modified Eagle’s medium (Mediatech Inc.), high glucose (4.5 g/L), supplemented with 10% fetal bovine serum (Hyclone) and 1 % Pen/Strep (Sigma). The cells were maintained in an incubator with 5% CO_2_ at 37 °C. HT1080 cells stably expressing RhoA FRET sensor was developed by Dr. Yi Wu at the University of Connecticut.

### Preparation of microfluidic devices

Molds to print the cell culture chamber and air chamber were fabricated by negative photoresist (SU8-2100, MicroChem Corp.). Typical soft lithography procedure was applied to fabricate our microfluidic devices. Briefly, a 200 µm-thick layer of SU8-2100 photoresist (MicroChem Corporation, Newton, MA, USA) was spun-coated onto a silicon wafer and cross-linked by UV light exposure through a photomask. Developer was used to remove non-crosslinked photoresist. Then 200μ*µ*m thick layer of PDMS (1:10 of agent to base, Sylgard 184, Dow Corning Corp.) was spun onto the mold of culture chamber and 7 mm thick PDMS was poured onto the mold of air chamber. Both layers with half-cured PDMS were carefully aligned and then baked until completely cured. Mold for micropillars of 4 *µ*m height was fabricated by patterning negative photoresist (SU8-2005, MicroChem Corp.) following similar procedures as described above. Then 100 *µ*m thick layer of PDMS was spun onto the mold and baked until fully cured. The PDMS layer with pillars was carefully peeled off the mold and assembled onto a coverglass, with pillars facing the air. The height of micropillars was measured by profilometer (Dektak IIA). Channel device containing both air chamber and cell culture chamber, 5-mm circular coverglasses and the micropillar PDMS on glass were treated with plasma system. The circular cover glass was then put in the middle of the air chamber. Finally, the channel device with the cover glass was assembled with the micropillar PDMS, with pillar side facing the air chamber.

Before the experiment, 0.2 ug/ml collagen I solution was added into culture chamber, left incubated at 37 °C for 1 hour for coating. Cells were then injected into culture chamber through tubing. After cells adhered and spread onto the collagen-coated PDMS surface, a moderate pressure (∼10 psi or 68 kPa) was applied through the tubing to the air chamber. The pressure was kept constant by a pressure regulator.

### Treatment of cells in compression device

After cells were perfused into the culture chamber of the device, they were allowed to attach and spread to the collagen-coated PDMS bottom for 2 hours. Then calcium free medium or normal cell culture medium with drug was perfused into the cell culture chamber through tubing. At least 3ml of medium was used to ensure the existing medium was all replaced. Cells were returned to incubator. After 10 to 30 minutes, the compression device containing cells was transferred to the live-cell unit mounted on the microscope, followed by imaging and compression. The concentration of TPRV4 inhibitor was 5 µM. The concentration of blebbistatin was 25 µM.

### Fixing and staining cells in the compression device

After cells were compressed for 30 minutes or overnight, the live-cell unit was turned off. PBS was perfused into the device slowly at room temperature. 3.7% PFA was then perfused in at 1ml/hr using a syringe pump for 30 minutes. Mechanical compression on the cells was released by removing air pressure. After permealizing cells using 0.1% Triton-X in PBS, primary antibody detecting YAP (1:100, Santa Cruz Techonology) was perfused into the device. Cells in the device was incubated with the primary antibody at 4°C overnight and then incubated with fluorescent secondary antibody for 2 hours at room temperature. Unbound antibodies were removed by continuously perfusing PBS through the device for 30 minutes at 3ml/hr using a syringe pump.

### Image Acquisition

All images were acquired with a Nikon TE2000E epifluorescence microscope (Nikon) equipped with Luca-R CCD camera (Andor Technology), an X-cite illuminator (Excelitas Technologies) and 40 x water immersion objective, NA=1.2, WD=200 µm (Nikon). Each cell was imaged using three scans: donor, FRET and acceptor, using the following band-pass filters: CFP (EX: 436/20, EM: 480/40); YFP (EX: 470/40, EM: 525/50); FRET (EX:436/20, EM:535/40). All images were taken at one-second exposure time. The pixel to length ratio is 1 pixel equals to 0.2 µm.

### Image Analysis and Data Acquisition

Since the pixel intensities within the cell region were much higher than the pixel intensities in the background region, a binary image based on the pixel intensities can be generated for each field of view, where pixels with high intensities in the cell region were marked with 1 and pixels elsewhere were marked with 0. We then used Matlab routine, “bwboundaries”, to trace the cell boundary from the binary image. Every traced region with area that was 1,500 pixels square (∼60 micrometer square) or lower was considered as debris or cell fragment, and, therefore, was ignored.

For HT1080 live cells stably expressing RhoA FRET sensor, we used the YFP channel to trace the cell boundary, because of its relatively stable fluorescence regardless whether the cells were compressed (Fig. S3). For fix and stain images, we used the YAP channel to trace the cell boundary and H2B channel to trace the nuclear boundary. The actual cell boundaries were then dilated 10 pixels away from the original boundaries, in order to capture all the scattered light from the epifluorescence source. The nuclear boundaries, however, are only dilated two pixels away from the traced nuclear boundary, in order to avoid excessive overestimation of nuclear YAP (Examples were shown in Fig. S2).

The total fluorescent intensity for one cell was computed by adding all the pixel intensities within a traced boundary, after subtracting the mean background intensity. We repeated each set of experiments three times for statistical analysis. For each experiment, we scaled each integrated fluorescent intensity by the mean integrated intensity of the uncompressed control counterpart.

## Results

### The air-driven microfluidic device to compress mammalian cells

To investigate the effects of mechanical compression on RhoA activity in mammalian cells, we designed an air-driven microfluidic compression device. Similar device was previously used to compress bacterial cells to investigate cell growth [27]. As illustrated in Fig. 1a, the device is comprised of an upper and a lower chamber separated by a PDMS layer of about 200 *um* in thickness. The upper chamber can be inflated by air pressure, which deforms the PDMS membrane downwards and applies mechanical compression on the mammalian cells cultured in the lower chamber. To precisely control the compression on the cells, micropillars were introduced to the device. Micropillars were made of PDMS and assembled onto the bottom of the lower chamber. The pillars support the PDMS membrane, and provide a maximum limit to the downward membrane movement, and thereby control the degree of compression of the mammalian cells. The height of the pillar is about 4 µm, which is smaller than the typical height of the adherent HT1080 cells on 2D substrates (about 8-15 µm). The size of the pillar is 500 µm by 500 µm, with 200 µm distance between them.

**Figure 1:**
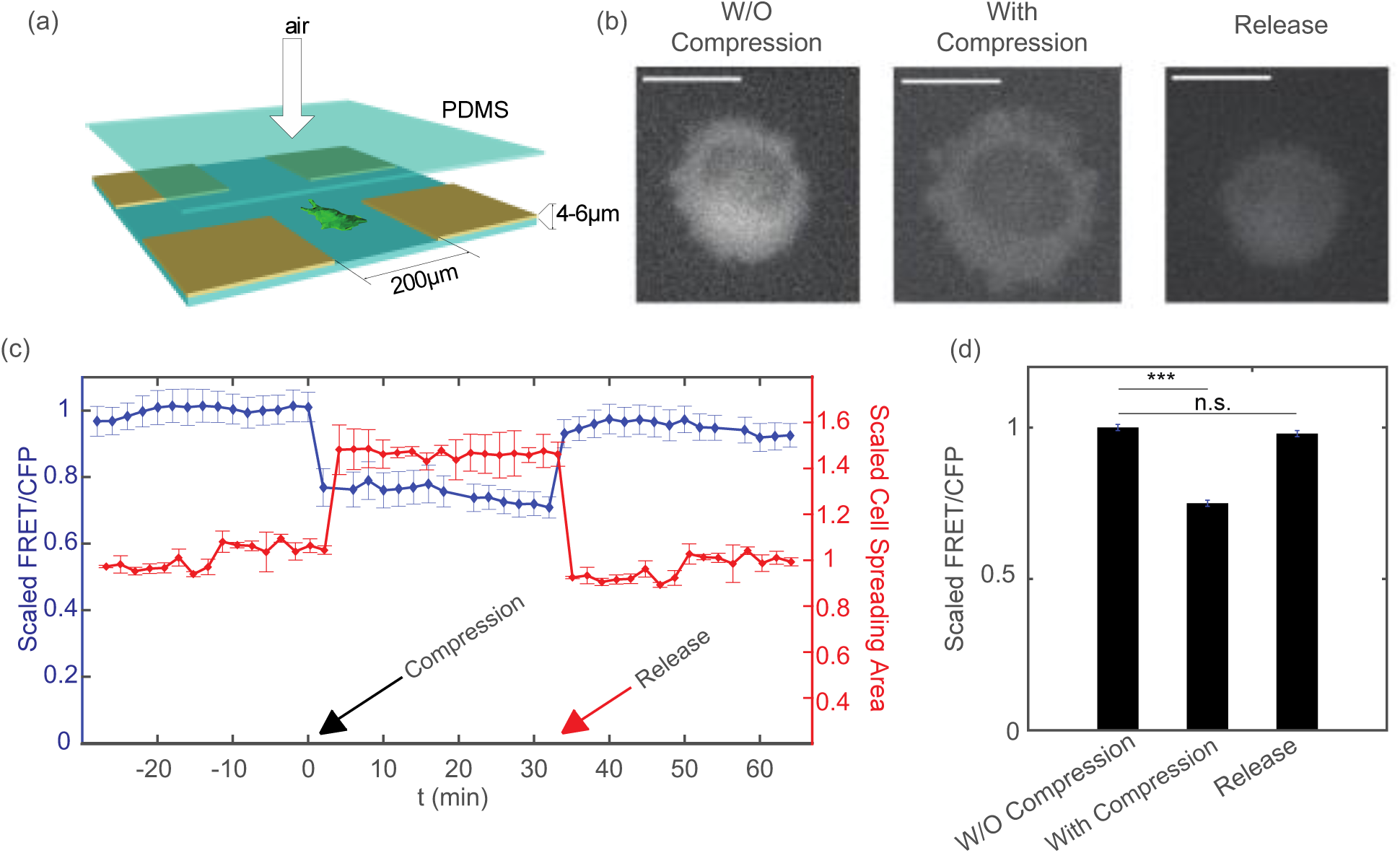
Response of HT1080 cells to mechanical compression. (a) Experimental setup for compression device. Fabrication of the device follows [27]. The cell in the device is compressed by the application of air pressure above the PDMS layer. **(b**) Epifluorescence image of cell (YFP channel) before and after compression, and after the release of compression. The cross-section area of the cell increases under compression, and decreases after the release of compression. **(c)** Scaled FRET/CFP and cell spreading area over time. Compression takes place at t = 0 min (Indicated by black arrow). The cells are then released at t = 38 min (indicated by red arrow) (n = 40 cells); **(d)** Summary of overall FRET/CFP when cell is uncompressed, compressed and released from compression. (All results are scaled with the mean value of uncompressed cells. Statistical significance: *** P<0.001. Scale Bar = 20 µm. n=40 cells).

To investigate the effects of mechanical compression on RhoA activity in mammalian cells, we used HT1080 cells stably expressing a RhoA FRET sensor. The intracellular sensor is comprised of two fluorescent proteins, RhoA GTPase, and a RhoA binding domain of the Rho effector PKN1 [28]. When RhoA-GDP is converted to RhoA-GTP, a PKN1 moiety binds RhoA-GTP, giving a high FRET state that is detected as an increase in sensitized emission over CFP ratio. We tested the sensor by applying a Rho activator and inhibitor. Results showed that the activity of RhoA sensor changed correspondingly as the activator or inhibitor was added to the cells (Fig. S1).

Before loading cells to the compression device, the cell culture chamber of the device was coated by 0.2 ug/ml collagen I to ensure the proper adherence and spreading of the cells onto the PDMS substrate. Then the cells were flown into the culture chamber through tubing. After cells adhered and spread onto the collagen-coated PDMS surface, a moderate pressure (∼10 psi or 68 kPa) was applied through the tubing to the air chamber. The pressure was kept constant by a pressure regulator. The downward movement of PDMS layer between the air chamber and the cell culture chamber stopped when the layer contacted micropillars, which applied a defined mechanical deformation on the HT1080 cells (Fig. 1a and b). During compression, a temperature of 37°°C was maintained and fresh medium was supplied by a constant flow.

### Instantaneous and reversible RhoA activity change upon compression

Previous work showed that RhoA activity increases and then decreases when cells are pulled by magnetic beads mechanically [10]. It is similarly interesting to probe the change of RhoA activity when live single cells are compressed vertically. We observed that the morphology of HT1080 cells changed instantaneously upon compression, as shown by a sudden increase in the observable size of the cell (Fig. 1b, middle panel, Fig. 1c). The ratio of the FRET channel and the donor CFP channel, which is an indicator of the RhoA activity, dropped significantly after the cells are compressed by the air-driven PDMS layer (Fig. 1c and d). The decrease in Rho activity is maintained constant throughout compression. To investigate whether such changes are reversible, we released the compression by turning off the air flow after 30 minutes of compression. We found that RhoA activity increased quickly back to the original value prior to compression, while there is also a simultaneous recovery in cell size (Fig.1 b-d). These results showed that changes in cell size and the RhoA activity are both instantaneous and reversible under compressive mechanical deformation. We also monitored and quantified the fluorescence intensity of YFP channel during the experiment, which should remain constant as long as the expression of RhoA sensor remains unchanged. Results showed that compression and release of compression did not affect the YFP intensity (Fig. S3), which further validates that changes in the ratio between the fluorescence intensities of FRET and CFP channel during compression and de-compression are not due to optical artifacts.

### Response of RhoA activity to mechanical compression is dependent on Calcium

In the carboxyl-terminal region of RhoA, there is a binding site for calmodulin, which is a ubiquitous transducer of calcium second messenger. A fusion protein was previously designed in which the activity of RhoA was controlled by calcium through calmodulin [29]. Moreover, previous research showed that Ca^2+^ influx is necessary for RhoA activation during human umbilical vein endothelial cell spreading on type IV collagen [30]. To investigate whether calcium is required for the response of RhoA activity to vertical compression, we incubated cells in calcium free medium after they attached and spread onto the substrate in the compression device.

After cells were incubated in calcium free culture medium for 10 minutes, they were subjected to compression within the device. We observed that cells in calcium free medium for 10 minutes had slightly reduced RhoA activity (Fig. 2b). After the cells were compressed, RhoA activity still was reduced significantly. However, the relative decrease in RhoA activity for cells incubated in calcium free medium was smaller compared to the change for cells incubated in normal culture medium (Fig. 2b and c). We also treated cells with calcium free medium for 30 minutes before compression. We find that longer incubation of cells in calcium free medium significantly decreased the activity of RhoA in cells without compression (Fig. 2b and c). The relative decrease in RhoA activity after compression was much smaller compared with cells in control medium and in calcium free medium for shorter incubation time (Fig. 2b). Taken together, our results of HT1080-RhoA cells in the compression device suggest that both the baseline RhoA activity and the response of RhoA to mechanical compression are calcium dependent.

**Figure 2:**
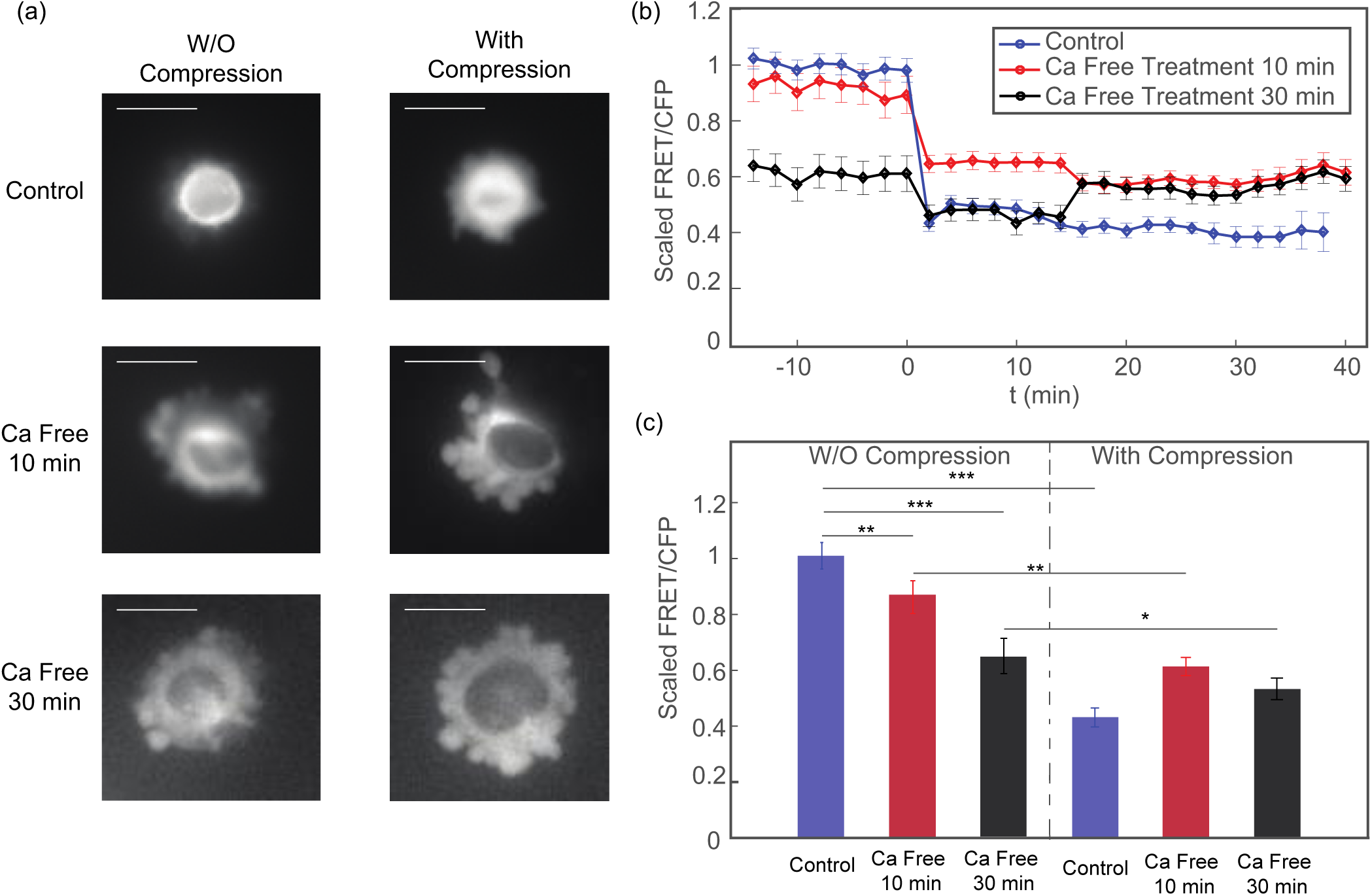
Response of cells to mechanical compression when they are incubated in Ca^2+^ free medium. Epifluorescence image (YFP channel) for cells before and after compression. When cells are incubated in Ca^2+^ Free medium, there is a significant increase in membrane blebs even without compression. **(b)** The scaled FRET/CFP ratio before and during the mechanical compression, for the three conditions in (a). Compression takes place at t = 0 min. The FRET/CFP ratio is scaled by the average FRET/CFP before compression for each respective control experiment. **(c)** Time averaged plots of FRET/CFP corresponding to panel (b). (Each set of results are scaled by its respective control, uncompressed mean FRET/CFP value. Statistical significance: ***P<0.0001; ** P< 0.001; *P<0.01. Scale Bar = 20 µm. n=50 cells for the control data, n=47 cells for 10min Ca free medium, n=27 cells for 30min Ca free medium.)

### TRPV4 channels mediate the change of RhoA activity during mechanical compression

We then set out to identify the mechanosenstive element that mediates the response of RhoA activity to mechanical compression. Recent studies showed that mechanosensitive membrane ion channels can regulate the activity of RhoA [31-33]. We are especially interested in transient receptor potential vanilloid 4 (TRPV4), a member of the TRP nonselective cation channel superfamily, because of its calcium permeability [34,35]. TRPV4 channels are expressed in both neuronal and non-neuronal cells, including HT1080 cells. Channel activation allows cation influx into cells, leading to various Ca^2+^ dependent processes. We have identified the role of calcium in the response of RhoA activity to vertical compression, which suggests the potential involvement of TRPV4 channel in regulating RhoA activity.

To investigate the role of TRPV4 in RhoA activity and the response to mechanical compression, we treated cells with a TRPV4 inhibitor before compressing them. We observed that incubating cells in TRPV4 inhibitor for 10 minutes did not affect RhoA activity before compression (Fig. 3b and c). Compressing the cells in the presence of TRPV4 inhibitor also lead to an instantaneous drop in RhoA activity, however, the magnitude of the change in RhoA activity due to compression was significantly smaller than that in control cells (Fig. 3b and c). RhoA activity in cells was reduced before compression after treated with TRPV4 inhibitor for a longer period of time, i.e. 30 minutes (Fig. 3b and c). There was essentially no change in RhoA activity after the cells are compressed in the presence of TRPV4 inhibitor. These results are consistent with those from the cells incubated in calcium free medium, suggesting that TRPV4 channels are indeed regulating the change of RhoA activity to mechanical compression through regulating calcium influx.

**Figure 3:**
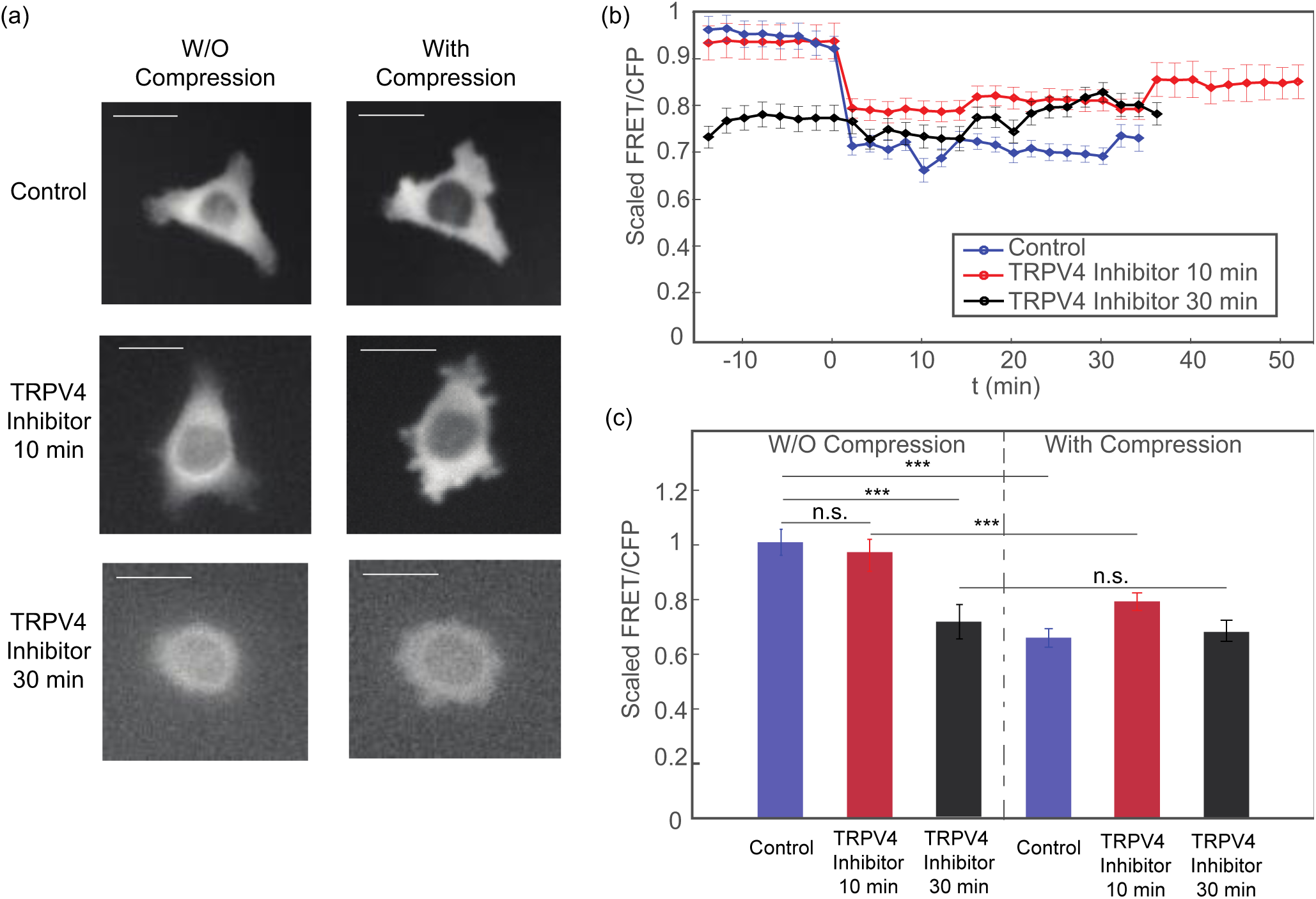
The response of cells to mechanical compression when they are incubated with TRPV4 inhibitor. (a) Epifluorescence image (YFP channel) for cells before and during compression. **(b)** The scaled FRET/CFP ratio before and during the mechanical compression, for all three conditions in (a). Compression takes place at t = 0 min. The FRET/CFP ratio is scaled by the average FRET/CFP before compression for each respective control experiment. **(c)** Time average plot of FRET/CFP corresponding to panel (b). (Each set of results are scaled by its respective control, uncompressed mean FRET/CFP value. Statistical significance: ***P<0.0001. Scale Bar = 20 µm. n =57 cells for the control data, n = 34 cells for 10 min TRPV4 inhibitor; and n = 30 cells 30 min TRPV4 inhibitor.)

### Mathematical Model of cell response to mechanical compression

The observed changes in Rho activation and the role of Ca flux can be recapitulated in a mathematical model of cell response to mechanical compression. When an external force, *F*_*ext*_, is applied to the cell, the applied force alters overall force balance at the cell surface (*F*_*ext*_ is negative if it is a compressive force). Mathematically, this force balance in the surface normal direction is expressed as [36,37]

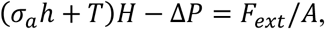

where *σ*_*a*_ is the active contractile stress in the cell cortex, *h* is the cortical thickness, *T* is the membrane tension, Δ*P* is the hydrostatic pressure difference across the membrane, *H* is the cell surface mean curvature, and A is the area over which the external force is applied. *F*_*ext*_ is the external mechanical force experienced by the cell, which is negative if it is a compressive force. Therefore, *F*_*ext*_ directly influences membrane tension, *T*, and results in the activation of TRPV4 and RhoA. Changes in active RhoA leads to changes in *σ*_*a*_, which re-adjusts membrane tension back to the homeostatic value. This feedback control loop can be expressed using kinetic equations as

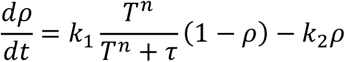

Where ρ is the proportion of active RhoA, and *k*_1,2_ are rate constants modeling activation and de-activation. The RhoA activation rate also depend on membrane tension and calcium through a Hill-function, and τ is an activation threshold parameter. This Hill-function explicitly describes the activity of TRPV4 as a function of membrane tension. As shown in Fig. 4(a) insert, the Hill function, Λ(*T*) = *T*^*n*^/(*T*^*n*^ + τ), can be approximated by a piecewise linear function. Activation threshold parameter, τ τ determines the critical membrane tension, *T*_*s*_, above which the channel is fully open. The active contraction is modeled as directly proportional to the amount of active RhoA: *σ*_*a*_(*t*) = Γρ(*t*), in which Γ is the contractile stress in the situation where RhoA is fully activated. This minimal model can approximately capture the behavior seen in experiments. Moreover, since mammalian cells can actively control the cytoplasmic pressure by adjusting their osmolyte and water content, water flows out of the cell when the hydrostatic pressure is increased, which reduces volume and membrane tension [36-38]. When active control of *σ*_*a*_ and Δ *P* are both incorporated the model is able to capture rate-dependent response of cells to external mechanical forces.

**Figure 4:**
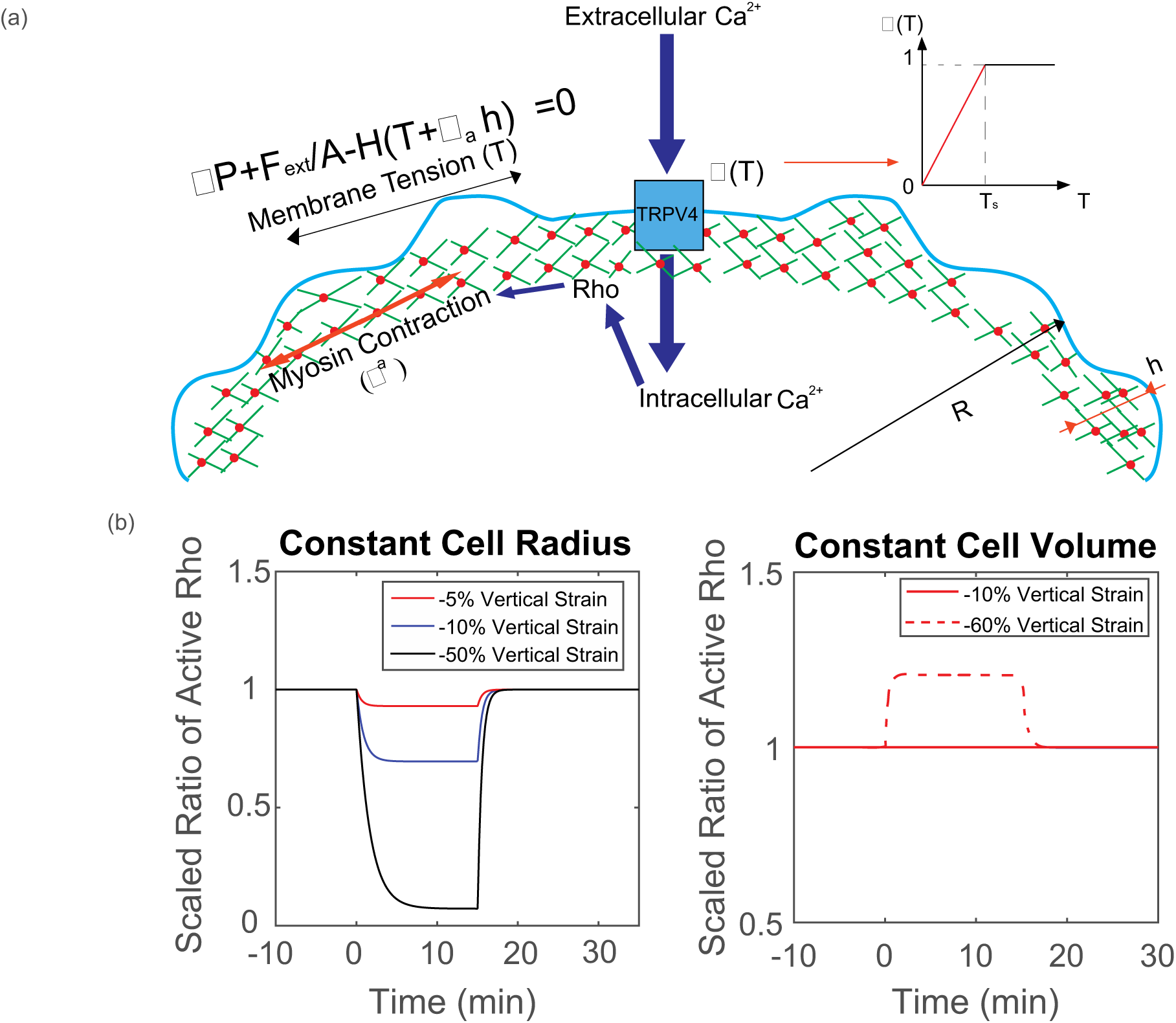
Theoretical prediction of cells under mechanical compression. (a) Mechanistic cartoon for our model: Membrane tension, *T*, and the active contractile stress from myosin contraction, *σ*^*a*^, are combined to balance the hydrostatic pressure difference across the cell membrane Δ*P*. Here, *H* is local mean curvature and *F*_*ext*_ is externally applied force, over the cross-section of the cell, *A*. Membrane tension changes activate Ca influx through TRPV4 channel, which influences Rho activity. The opening probability of TRPV4 channel is described by Hill function, which can be approximated by piecewise linear function; Model prediction of RhoA activity in a cylindrical cell in response to vertical compression when the cell maintains a constant cross-sectional area and constant volume. Depending on the extent of vertical compression, reduction of RhoA activity is observed when the cell reduces volume. If the volume remains constant, RhoA activity generally increases. In the experiment, there is a wide range of vertical compression. The observed average RhoA activity decrease suggests that there is a slight reduction in cell volume.

Based on this model, we can examine two extreme scenarios when the cell is under mechanical compression: cells that maintain a constant cross sectional area and cells that maintain a constant volume (i.e. constantΔ*P*, no water flow across the membrane). When cells maintain a constant cross sectional area, the lateral membrane tension decreases as the volume of the cell decreases. In addition, apical membrane surface of the cell in contact with the compression surface will experience lower tension. These factors combined suggest that RhoA activity goes down. When cells maintaining a constant volume, lateral membrane tension increases as the cell is compressed in the vertical direction. Therefore, RhoA activity would increase in the lateral surfaces. The overall measured RhoA activity would increase.

In Fig. 4, we compare and contrast these 2 scenarios for a model cylindrical cell under compression. Different degrees of compression and computed RhoA activities are plotted as functions time. We find that the experimental results are best explained by the case where water can flow out of the cell. In the SM we also use the same model to compute the cell response for a more realistic cell geometry and find a similar result. We can conclude that when the cell is under vertical compression, the behavior is somewhere in between the model extremes discussed above: the overall cell volume decreased, despite the fact that the cell cross sectional area increased, leading to a lower membrane tension and lower RhoA activity.

### Change of YAP subcellular localization upon mechanical compression

Recent research revealed the significant role that YAP (Yes-associated protein) plays in relaying the mechanical signal in extracellular matrix (ECM) to nucleus [14]. Previous work also showed that the subcellular location of YAP is regulated by cell tension and cell geometry, i.e. the cell spreading imposed by ECM [14,39,40]. In our compression device, we observed that the observed area of the compressed cells changes significantly in morphology. Thus, we hypothesized that the subcellular localization of YAP also changes upon mechanical compression.

To test our hypothesis, we fixed the compressed HT1080 cells 30 minutes or 13 hours after subjecting the cells to compression, and then we permeabilized the cells and stained them for YAP. Cells in uncompressed area showed predominant expression of YAP in cell nucleus (Fig. 5a, top panel). On the other hand, cells under compression showed much more signal of YAP staining in the cytoplasm (Fig. 5a, bottom panel). With the help of H2B-mCherry, a fluorescence labeled histone protein, we were able to quantify the amount of YAP staining in the nucleus and in the cytoplasm. Results showed that there was a dramatic and quick change in YAP subcellular localization upon mechanical compression. The ratio between YAP in nucleus and in cytoplasmic decreased by more than 60% after compression, i.e. most of the YAP “leaked out” the nucleus. There is no difference in cells compressed for 30 minutes or 13 hours. Interestingly, we also find that, while YAP “leaks out” of the nucleus when cells are compressed, cells are expressing more overall YAP, as shown in Fig. S5.

**Figure 5:**
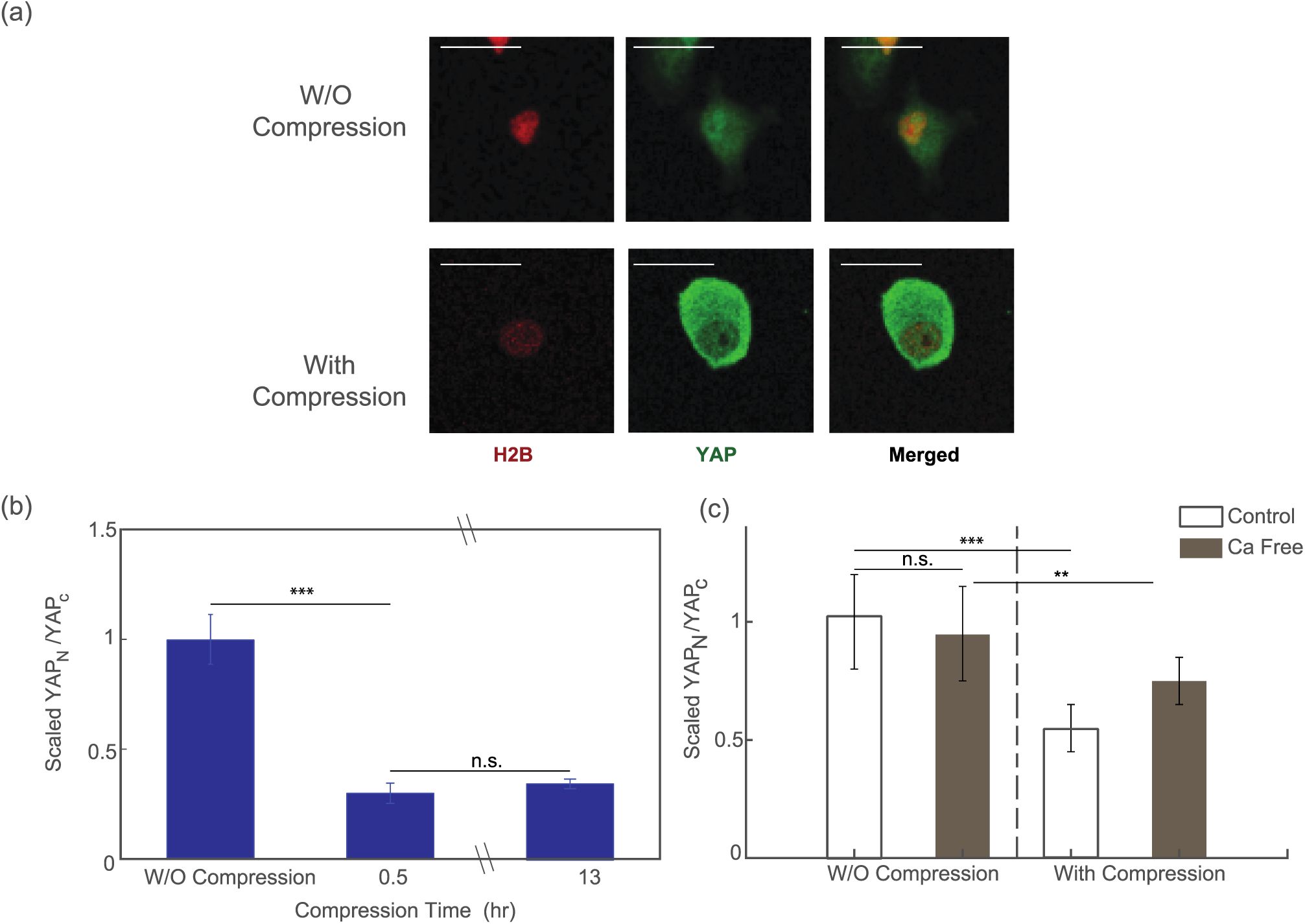
YAP expression level before and after compression. (a) Fix and stain images of the cells before (top panel) and after (bottom panel) mechanical compression. **(b)** Ratio between the nucleus YAP and cytoplasmic YAP before and after compression. Around 70% of nucleus YAP leaked out into cytoplasm after compression. The equilibrium is reached at around 30 minutes after compression takes place. (n=60 cells before compression, n=70 cells for post-compression after 30 min, n=84 cells for post-compression after 13 hours.) **(c)** The change in the localization of YFP during mechanical compression reduces in the Ca^2+^ deficit condition, such that the mechanosensation of YAP is suppressed. (n = 23 cells for control, pre-compression data, n = 30 cells for control, post-compression data. n = 27 cells for Ca free, pre-compression data and n = 36 for Ca free, post-compression data.) Statistical significance: ***P<0.0001; **P<0.01, *P<0.05.

We further investigated the response of YAP localization in calcium free medium. Results showed that export of YAP from the nucleus to the cytoplasm during compression was suppressed when the cells were deprived of calcium, suggesting that the translocation of YAP during mechanical compression is dependent on calcium (Fig. 5(c)).

## Discussion and Conclusions

Physical environment of the cell can influence many aspects of cell function, and cells have developed sensory systems to respond to environmental changes. By examining cells under mechanical compression, we discover that the activity of RhoA changes quickly in response to the externally applied force. Under compression, RhoA activity decreases, and recovers if the compression is removed. We found that RhoA activity is regulated by Ca^2+^ and mechanosensitive cation channel TRPV4. When TRPV4 is blocked, RhoA activity decreased before mechanical compression, and the response to compression is also less pronounced. Since RhoA is directly involved in phosphorylation of myosin light-chain and generation of active contractile stress, these results implicate Ca^2+^ as a major regulator of cellular mechanosensation, in agreement with evidence from other lines of investigation [24, 41]. Moreover, these results suggest that the cell membrane and associated channels are major elements in mechanosensation, in agreement with previous suggestions [42,43].

The results also suggest that myosin contraction is involved in regulating cell membrane tension. This is consistent with a theoretical model of the cell cortex where cortical contraction and membrane tension are together balancing excess hydrostatic pressure in the cell. The excess hydrostatic pressure, which is on the order of 100-1000Pa, is generated from excess osmotic pressure inside the cell. Most of this excess pressure is balanced by cortical contraction, and membrane tension is maintained at a low value. During rapid changes in osmolarity or externally applied mechanical force, the membrane tension changes rapidly, which changes the Ca^2+^ currents and myosin contraction, so that the membrane tension can be restored back to the homeostatic value. Therefore, the cortex acts as an active mechanical surface that dynamically adjusts its tension to changing external osmotic or mechanical perturbations. The active adjustment of cortical contraction is mediated by RhoA activity, signaled by Ca^2+^ currents. The theoretical model of the cell cortex can fully explain the observed data, and predicts many features of active responses of cells to external forces and varying stiffnesses of the cell substrate.

If we attempt to alter myosin activity through non-calcium related pathways, for example by incubating cells in 25uM blebbistatin for 1 hour, our experimental results show that, on average, blebbistatin only decreases RhoA activity slightly, and the relative drop in RhoA activity due to mechanical compression is similar to that of control experiment (Fig. S4). This means that blebbistatin, although suppresses myosin contractility, does very little to influence mechanosensitivity of RhoA that is upstream to myosin. We also find that mechanical compression also directly impact YAP transcription nuclear localization on the time scale of hours. YAP has been identified as a mechanosensitive transcription factor directly involved in mechanotransduction. Our results are consistent with the idea that YAP is sensitive to mechanical state of the cell. In addition, our data reveal that RhoA and myosin contraction should change rapidly (∼mins) in response to changes in mechanical force. Therefore, YAP activity may be directly regulated by Rho or myosin activity.

As an additional observation, we find that the mechanosensitivity of YAP may also depend on the cell shape, as more rounded cells have more YAP leaking out of nucleus compared to more elongated cells (Fig. S6). However, this may be due to the fact that more rounded cells are taller, and, therefore, are more compressed than the elongated cells.

Taken together, our mechanical compression experiments and model reveal that cells sense mechanical changes from changing membrane tension and calcium currents, and dynamically adjust contractile forces to balance external forces. The active regulation of cell contractile forces can also serve as a signal to change transcription factor localization and protein expression changes that completes the process of mechanotransduction.

## Acknowledgements

The present work was has been supported by NIH grant R01GM114675 and U54CA210172.

## Supplemental Figure legend

**Figure S1:**
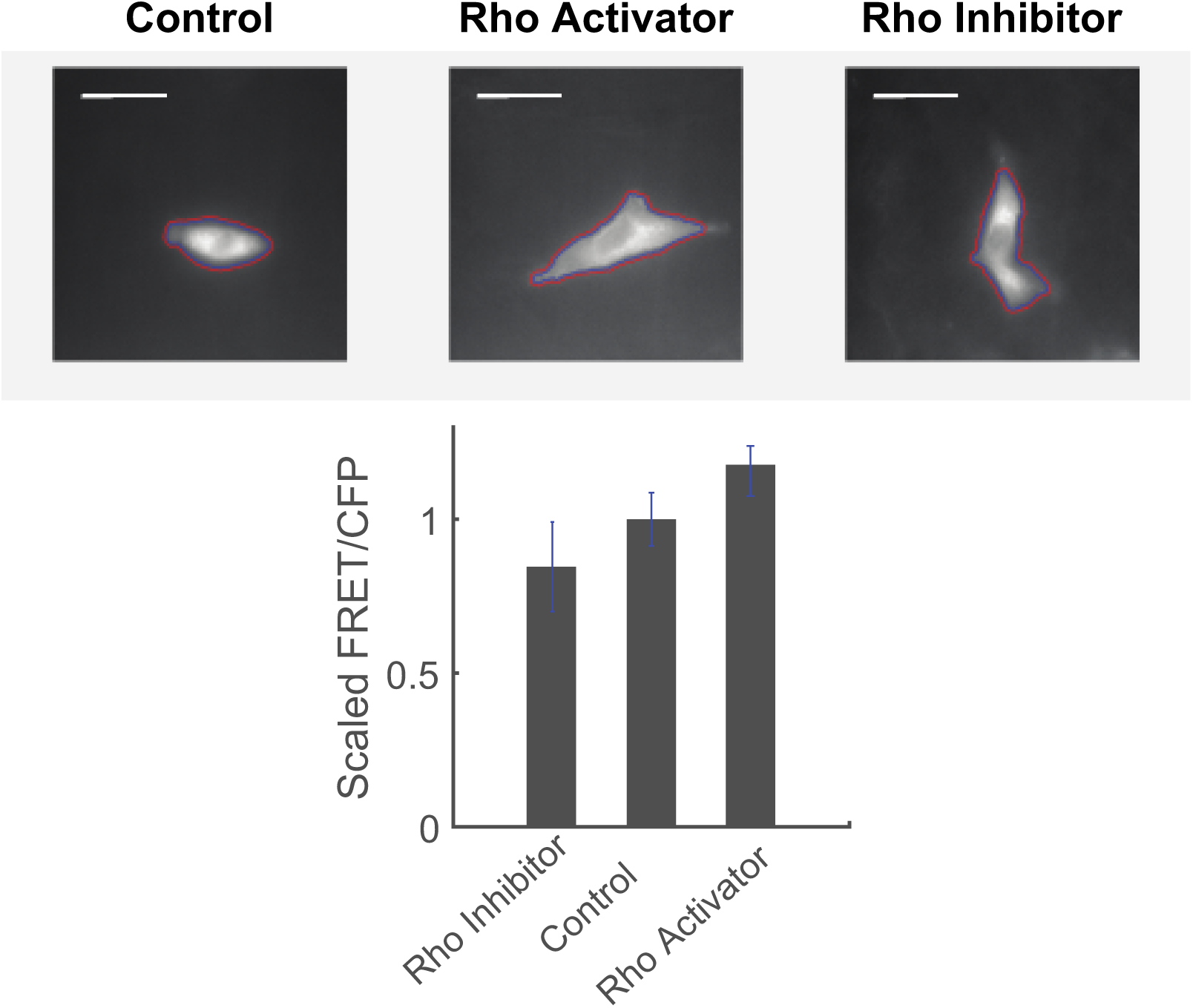
HT1080 cells stably expressing RhoA FRET sensor were treated with 1 unit/ml Rho activator (Cytoskeleton, CN01) for 30 minutes or 2 µg/ml inhibitor (Cytoskeleton, CT04) for 4 hours, respectively. RhoA inhibitor and activator significantly increases or decreases the activity of the RhoA sensor after the respective treatment. Scale bar = 20 µm.

**Figure S2:**
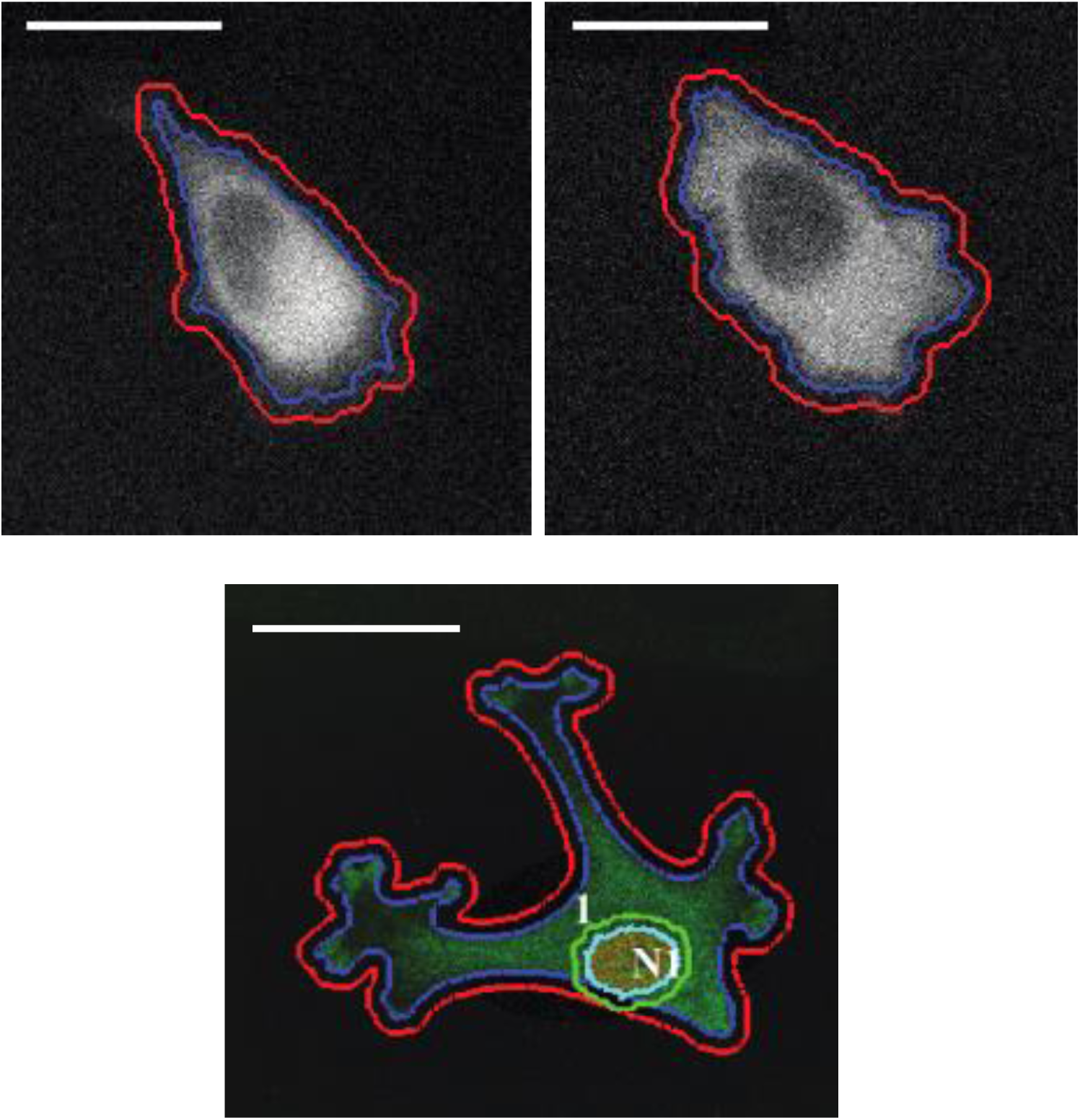
Tracing Examples of fluorescence images. (a) Examples of HT1080 live cells from YFP channel. **(b)** Examples from fixing and staining. Scale bar = 20 µm.

**Figure S3:**
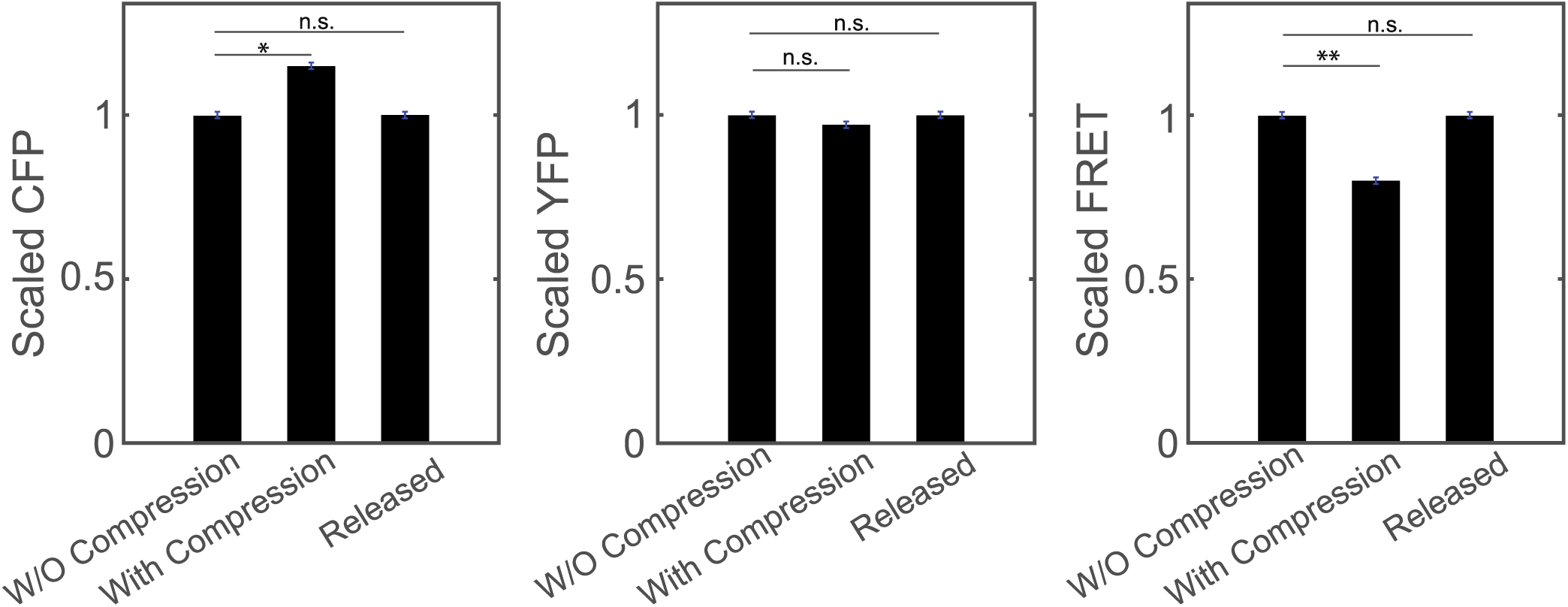
Change in three fluorescence channels when cell is under compression. The fluorescence intensities are scaled with pre-compression intensity value. The overall CFP intensity increased by 10 to 15% when compression is applied; overall YFP intensity remains more or less a constant (within 1% variation); while FRET intensity decreases about 15 to 20% when compression took place. Statistical significance: *P<0.05; ** P<0.01.

**Figure S4:**
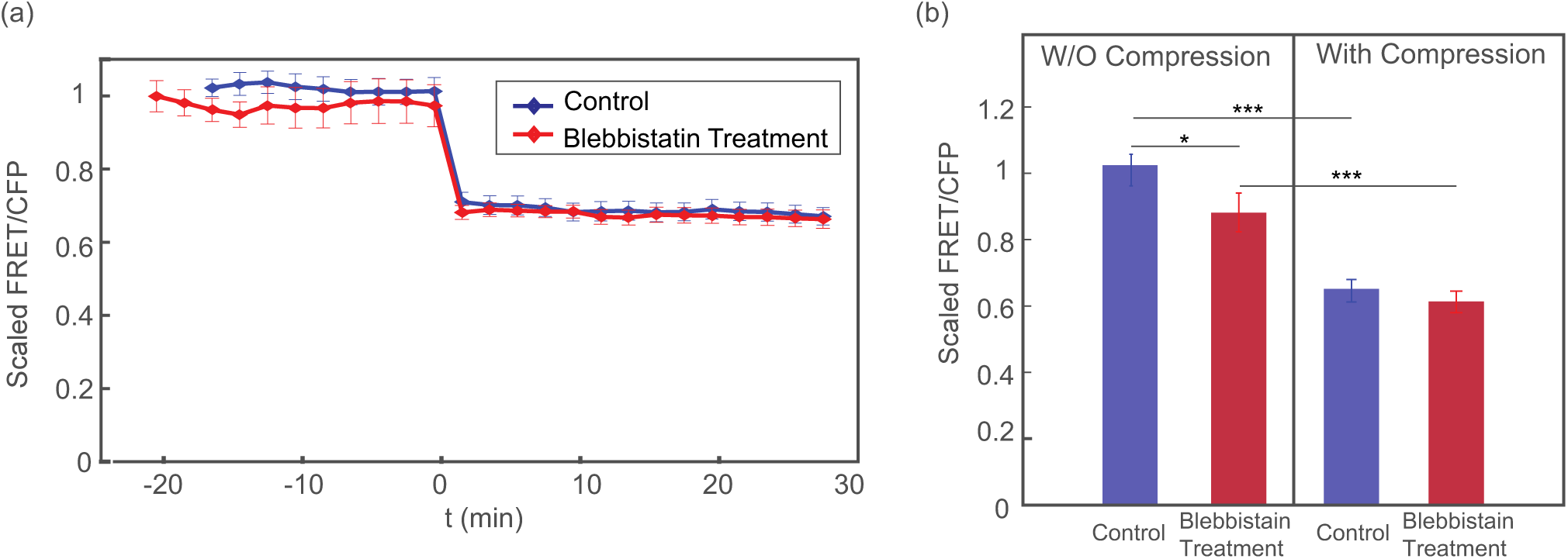
Blebbistatin treatment does not significantly affect mechanosensation of RhoA when cells are subject to mechanical compression. HT1080 cells stably expressing RhoA FRET sensor was incubated in 25 uM blebbistatin for one hour before the imaging took place. **(a)** Overall FRET/CFP ratio over time. Compression takes place at t = 0 min. **(b)** Time average FRET/CFP in terms of mechanical compression. Before compression, blebbistatin does slightly decrease overall RhoA activity, but it does not affect the change in RhoA when compression takes place. (Statistical significance: *P <0.01*** P<0.000001. n = 34 cells for control data and n = 28 cells for blebbistatin data.)

**Figure S5:**
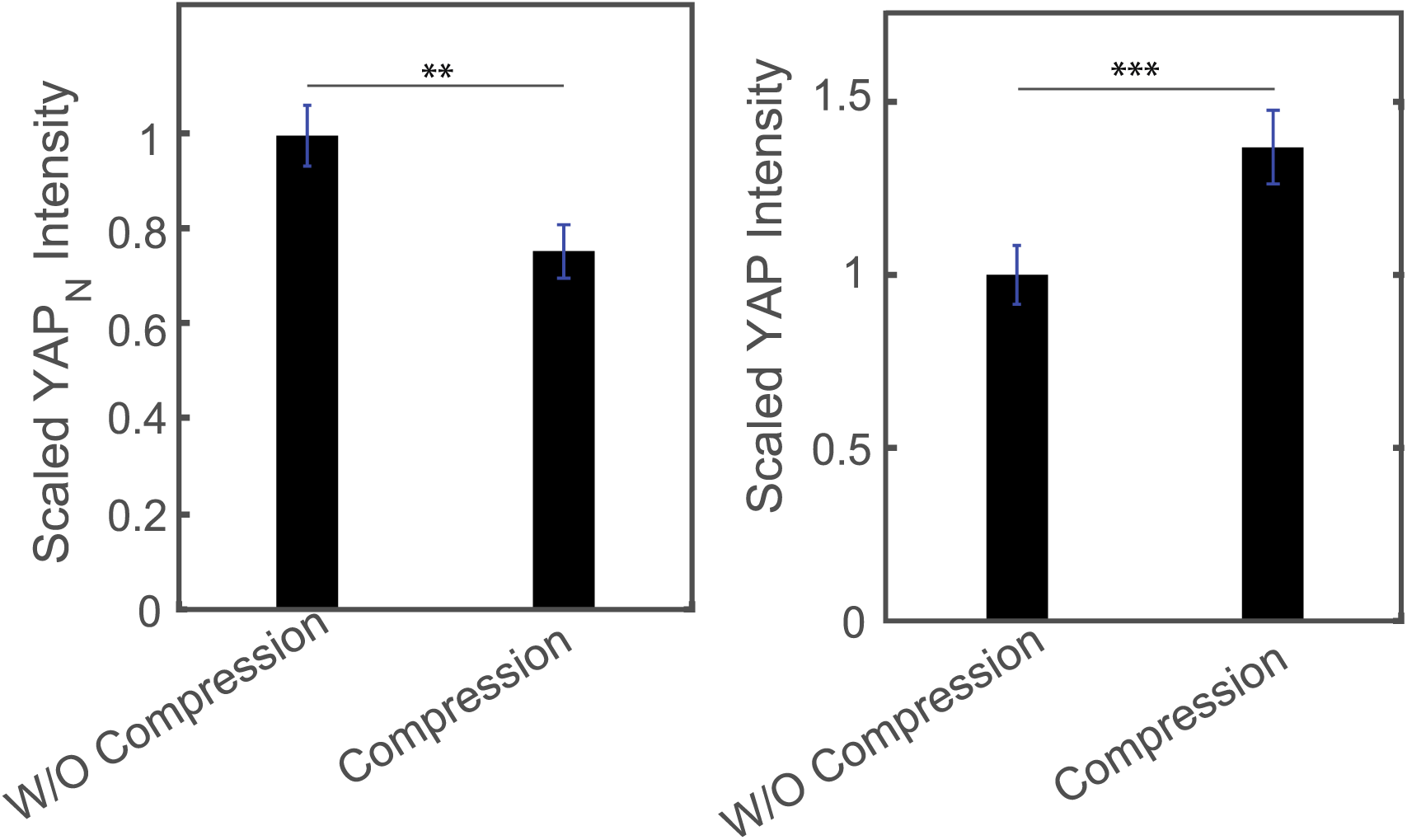
Reports of YAP Intensity of cells with and without compression. When cell is under compression, cell nucleaus YAP decreases, which has the consistent trend of Rho activity **(a)**. However, compressed cells also seem to produce significantly more total YAP than uncompressed cells **(b).** Statistical Significance: ** p<0.01, *** P<0.001.

**Figure S6:**
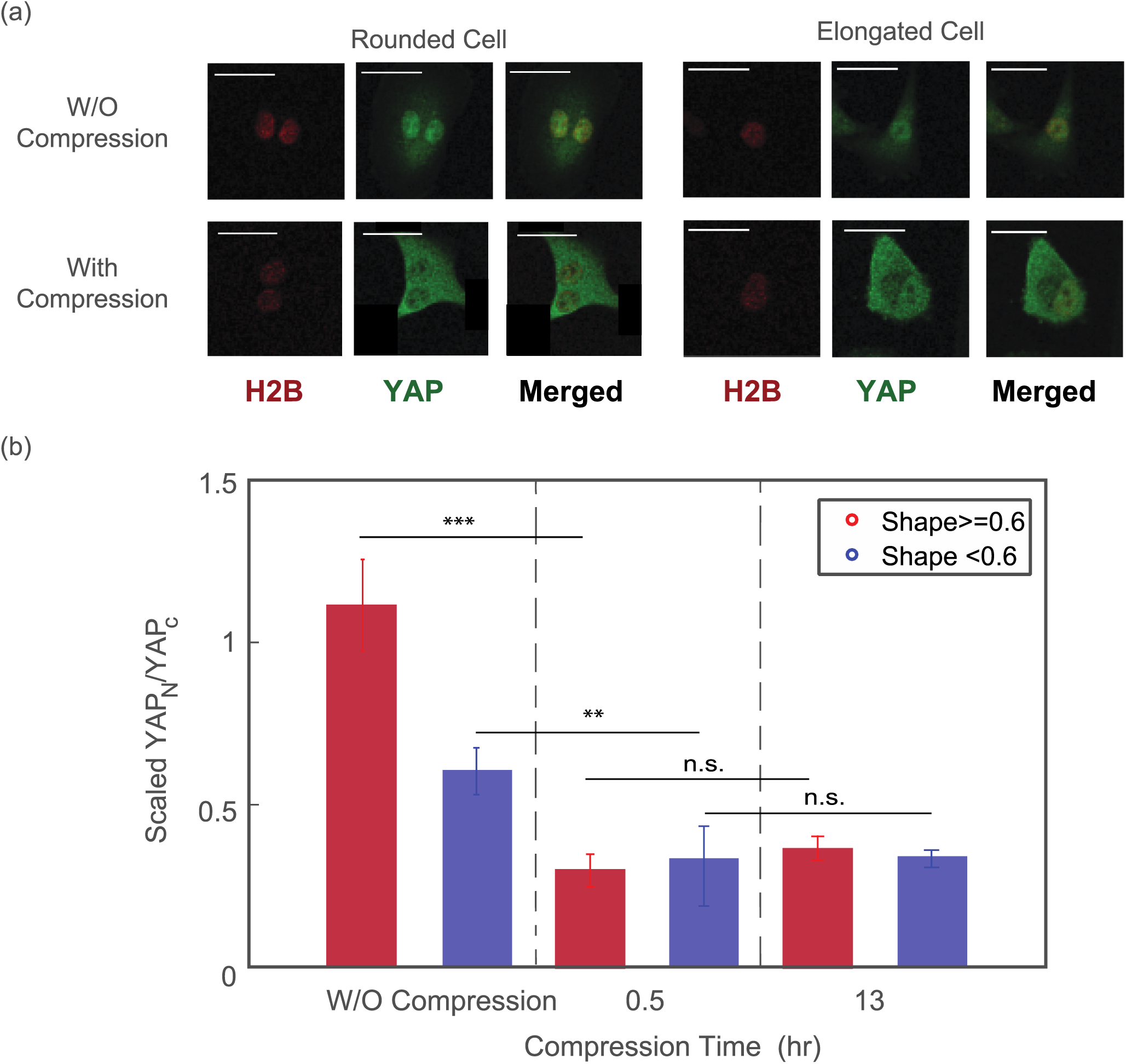
Mechanosensation of YAP may depend on the cell shape. (a) Examples of rounded cell and elongated cell before and after compression; **(b)** Ratio of Nucleus YAP in response of mechanical compression. For the elongated cells, YAP is less sensitive to mechanical compression. Statistical significance: ***P<0.0001; **P<0.01; Scale Bar = 20 μm. Here, the cell shape factor is defined as: 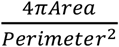; cell is more rounded if the shape factor is close to 1, and is more elongated if the shape factor goes to zero.

**Figure S7: Model calculation with realistic cell geometry. (a)** Schematic description of cell geometry before and after compression; (b) Corresponding RhoA activity, overall membrane area, membrane tension and cell volume in response to vertical compression. With a realistic cell geometry, results are similar to predictions from the simplified model described in the main text.

